# Transformations of sensory information in the brain reflect a changing definition of optimality

**DOI:** 10.1101/2023.03.24.534044

**Authors:** Tyler S. Manning, Emma Alexander, Bruce G. Cumming, Gregory C. DeAngelis, Xin Huang, Emily A. Cooper

## Abstract

Neurons throughout the brain modulate their firing rate lawfully in response to changes in sensory input. Theories of neural computation posit that these modulations reflect the outcome of a constrained optimization: neurons aim to efficiently and robustly represent sensory information under resource limitations. Our understanding of how this optimization varies across the brain, however, is still in its infancy. Here, we show that neural responses transform along the dorsal stream of the visual system in a manner consistent with a transition from optimizing for information preservation to optimizing for perceptual discrimination. Focusing on binocular disparity – the slight differences in how objects project to the two eyes – we re-analyze measurements from neurons characterizing tuning curves in macaque monkey brain regions V1, V2, and MT, and compare these to measurements of the natural visual statistics of binocular disparity. The changes in tuning curve characteristics are computationally consistent with a shift in optimization goals from maximizing the information encoded about naturally occurring binocular disparities to maximizing the ability to support fine disparity discrimination. We find that a change towards tuning curves preferring larger disparities is a key driver of this shift. These results provide new insight into previously-identified differences between disparity-selective regions of cortex and suggest these differences play an important role in supporting visually-guided behavior. Our findings support a key re-framing of optimal coding in regions of the brain that contain sensory information, emphasizing the need to consider not just information preservation and neural resources, but also relevance to behavior.

**Significance:** A major role of the brain is to transform information from the sensory organs into signals that can be used to guide behavior. Neural activity is noisy and can consume large amount of energy, so sensory neurons must optimize their information processing so as to limit energy consumption while maintaining key behaviorally-relevant information. In this report, we re-examine classically-defined brain areas in the visual processing hierarchy, and ask whether neurons in these areas vary lawfully in how they represent sensory information. Our results suggest that neurons in these brain areas shift from being an optimal conduit of sensory information to optimally supporting perceptual discrimination during natural tasks.

---

An appealing theory of neural computation is that neurons in early sensory areas respond to stimuli in a way that maximizes the information carried about the world (1, 2) while neurons in downstream areas transform this representation so as to best support specific tasks and computations (3–6). A careful test of this theory of sensory transformations requires several elements: we need to characterize the typical probability distribution of a pertinent sensory variable encountered in the environment, we need large-scale measurements of neural responses driven by this sensory variable across multiple brain areas, and we need a single mathematical framework that can be applied to sensory representations shaped by different behavioral or computational objectives. Here, we use binocular disparity in the primate visual system as a model, and combine across diverse data sets to test this theory. Our results provide strong empirical evidence in support of a systematic transformation of sensory representations in the brain: from information preservation at early processing stages to maximizing perceptual discrimination performance at later stages.

Binocular disparity between the retinal images offers an ideal test bed for examining hierarchical sensory representations. In animals with forward-facing eyes, non-fixated points in space tend to fall on disparate retinotopic locations because the eyes are laterally offset from each other (Fig. 1A). Successful integration of information from the two eyes relies on populations of neurons that are tuned for different binocular disparities–the differences in the retinotopic location of images in the left and right eyes. While neuronal tuning for binocular disparity emerges early in the mammalian visual system (V1), populations of neurons tuned for binocular disparity have also been characterized all along the dorsal and ventral processing streams (7, 8). Beyond just binocular integration, sensing of horizontal binocular disparities in particular provides one of the most reliable cues to the relative distances of objects in the environment, and as such this cue supports a variety of high-level perceptual tasks such as figure/ground segmentation, 3D motion perception, and breaking camouflage (9–11). Indeed, the magnitude and direction of horizontal binocular disparity varies lawfully as a function of how far objects are from the observer as well as where the observer is fixating, and prior work has shown that these variations result in predictable statistical regularities in the binocular disparities encountered during natural tasks (12–15). We hypothesized that early representations of horizontal binocular disparity maximize the information carried about typical disparities encountered during natural behavior, while later representations instead facilitate discrimination of disparity to support specific perceptual tasks.

**Fig. 1.**
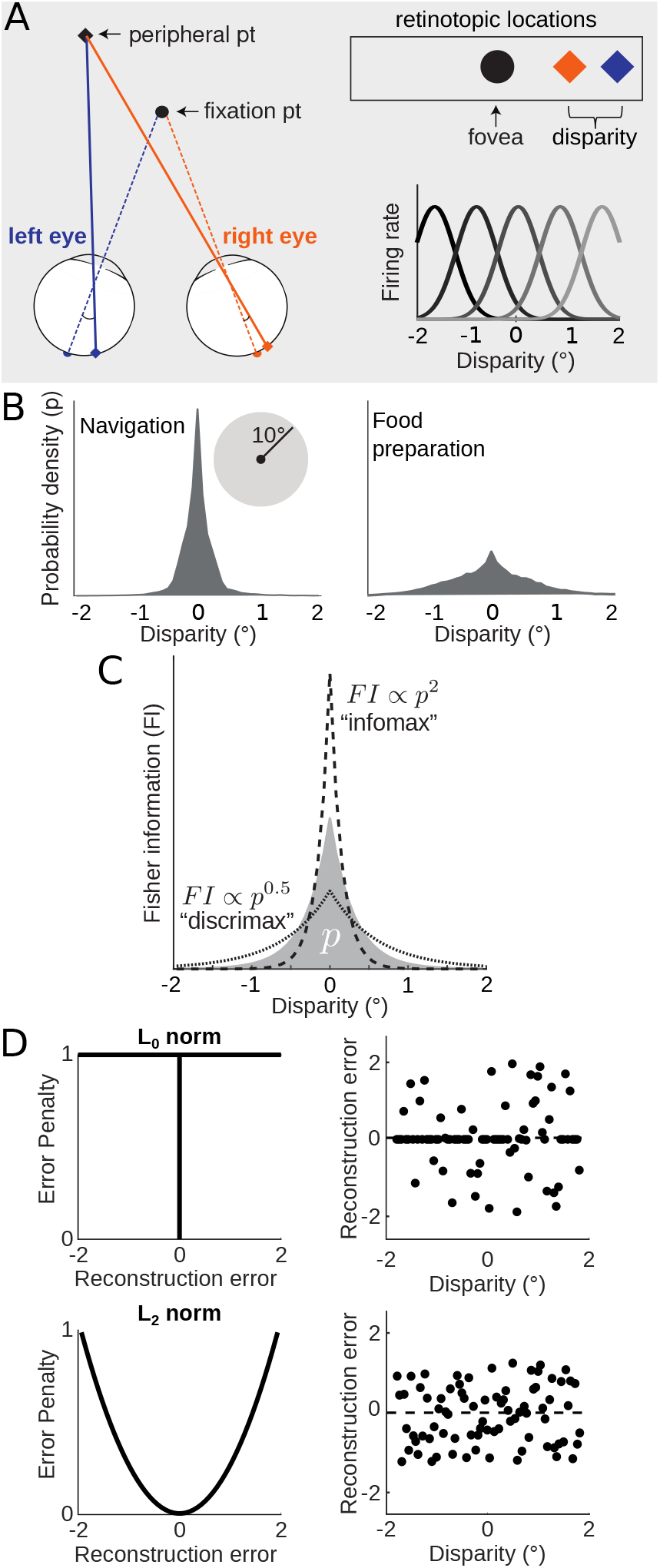
**A.** Points in the peripheral visual field tend to fall on disparate retinotopic locations because the eyes are laterally offset. The retinotopic difference in these locations is called the horizontal binocular disparity (abbreviated as simply disparity in figures). Populations of neurons that are tuned for different horizontal disparities, illustrated in the bottom right, are found throughout the visual system. **B.** Disparities encountered in the central visual field (10° of fixation) tend to be small. Each plot shows the probability density distribution of binocular disparity obtained from data collected by (12) while human participants either navigated an indoor environment (Left) or prepared a sandwich (Right). **C.** Information theoretic frameworks indicate that optimal sensory representations can be described by a power law relationship between disparity stimulus probability *p* (shaded region) and the Fisher Information (*FI*) of a neuronal population. Information maximizing codes (“infomax”, dashed line) are proportionate to the squared probability, while discrimination maximizing codes (“discrimax”, dotted line) are characterized by a compressive nonlinearity. **D.** The plots on the left illustrate normalized error functions that minimize the *L*_0_ (top) and *L*2 (bottom) norms. The plots on the right illustrate example patterns of disparity estimate errors under each of the respective norms used to optimize the reconstruction of the ground truth from noisy sensory measurements. For clarity, disparities are sampled from a uniform distribution. The sets of reconstruction error under the *L*_0_ and *L*2 norm have equal total error penalty. Dashed line indicates zero error.

## Results

### Natural distributions of binocular disparity have strong statistical regularities

To test this sensory transformation hypothesis, we first need an understanding of the distribution of horizontal binocular disparities that the visual system is tasked with processing (hereafter simply referred to as binocular disparities). In recent years, there has been a concerted effort to characterize the visual “diet” of binocular disparities that is typical of natural experience (12–15). This work suggests several robust statistical properties of typical binocular disparities, most notably that small disparities (near zero) tend to be much more likely than larger disparities. Optimal sensory representations for binocular integration should therefore differentially allocate processing resources for small disparities versus large ones. Using a previously-collected data set in which people were eye-tracked while performing natural tasks (12), we calculated the probability distribution of binocular disparities within a central 10° radius from fixation while people performed two different tasks: food preparation and navigation (Fig. 1B). While the precise distribution shape differed between the tasks, we see that both distributions are approximately zero-mean, symmetric, and highly kurtotic (consistent with previous analyses). The increased prevalence of larger disparities during food preparation is likely due to the different typical object distances between the tasks: a given depth interval between two objects maps to a larger disparity if the objects are relatively close to the observer, as during manual tasks like preparing food.

### Sensory coding theories predict a lawful transition in how these binocular disparity statistics are reflected in the brain

Optimal coding frameworks provide concrete, testable predictions for how a stimulus probability distribution should be reflected in neural populations. Specifically, if a population maximizes the information carried about a sensory variable, we often expect the Fisher Information of the population neural activity *(FI*; a measure of the precision with which a sensory variable is encoded) to follow a power law in which it is proportionate to the stimulus probability squared (Fig. 1C, “infomax” dashed line)(16, 17). This representation can be interpreted as a reference prior (18), which assumes the least possible information about the world, or Jeffrey’s prior, providing invariance across transformation of sensory units (19).

Under common assumptions on computational limits, this *FI* ∝ *p*^2^ power law corresponds to a neural code that minimizes the *L*_0_ norm of the stimulus reconstruction error (5, 6, 16, 17). This means that all error magnitudes are penalized equally and the expected value of these errors is minimized (i.e., error sparsity is maximized; Fig. 1D, top) (5, 6). Prior work has found signatures of this power law across a range of early sensory brain areas (3, 20). However, a representation with this distribution is not optimal for visual tasks that require discrimination between different values of a stimulus variable, because error magnitude matters for many tasks. For example, if one is trying to determine whether their hand can fit in between two sharp objects separated in depth, a small error may be harmless but a large error may lead to injury. Thus, for perceptual discrimination, neural codes that minimize some other error metric like mean squared error are often appropriate (*L*_2_ norm; Fig. 1D, bottom). As such, codes that are optimized for downstream discrimination tasks should reduce the concentration of neural processing resources on high probability events and spread *FI* more equally across the stimulus space (Fig. 1C, “discrimax” dotted line)(5, 17). The discrimination-maximizing “discrimax” line shown in the Figure corresponds to a specific power law of *FI* ∝ *p*^0^.5. This objective (along with objects that aim to minimize other *L_p_* norm errors like the sum of absolute errors) results in consistently more compressive nonlinearities than information maximization (6, 21)). For neural codes for binocular disparity, we would therefore expect the population *FI* to be more strongly peaked at zero disparity in early visual regions, and more equally spread out across a broader range of disparities in later visual regions.

### Neural populations differ as predicted by a transition from information-preservation to supporting perceptual discriminations

To test this transformation hypothesis, we must characterize a large number of neuronal tuning curves for binocular disparity across different brain areas, such that we can calculate the *FI* associated with these tunings and compare them to the disparity probability distributions in 1B. Since the precise shape of the disparity probability distribution varies between tasks (and different resource constraints can change the numerical value of the optimal exponent for the power law (5, 21)), here we focus on the *relative* transformation of the *FI* exponent between brain regions rather than on its nominal value. To this end, we compiled a data set of 1056 neurons’ binocular disparity responses spanning brain areas V1, V2, and MT of the macaque monkey. The mean responses of each neuron as a function of binocular disparity were fit with a continuous 1D Gabor function and the individual neuron *FI* associated with each tuning curve was calculated from these fits based on assumption of Poisson spiking (Fig. 2A).

**Fig. 2.**
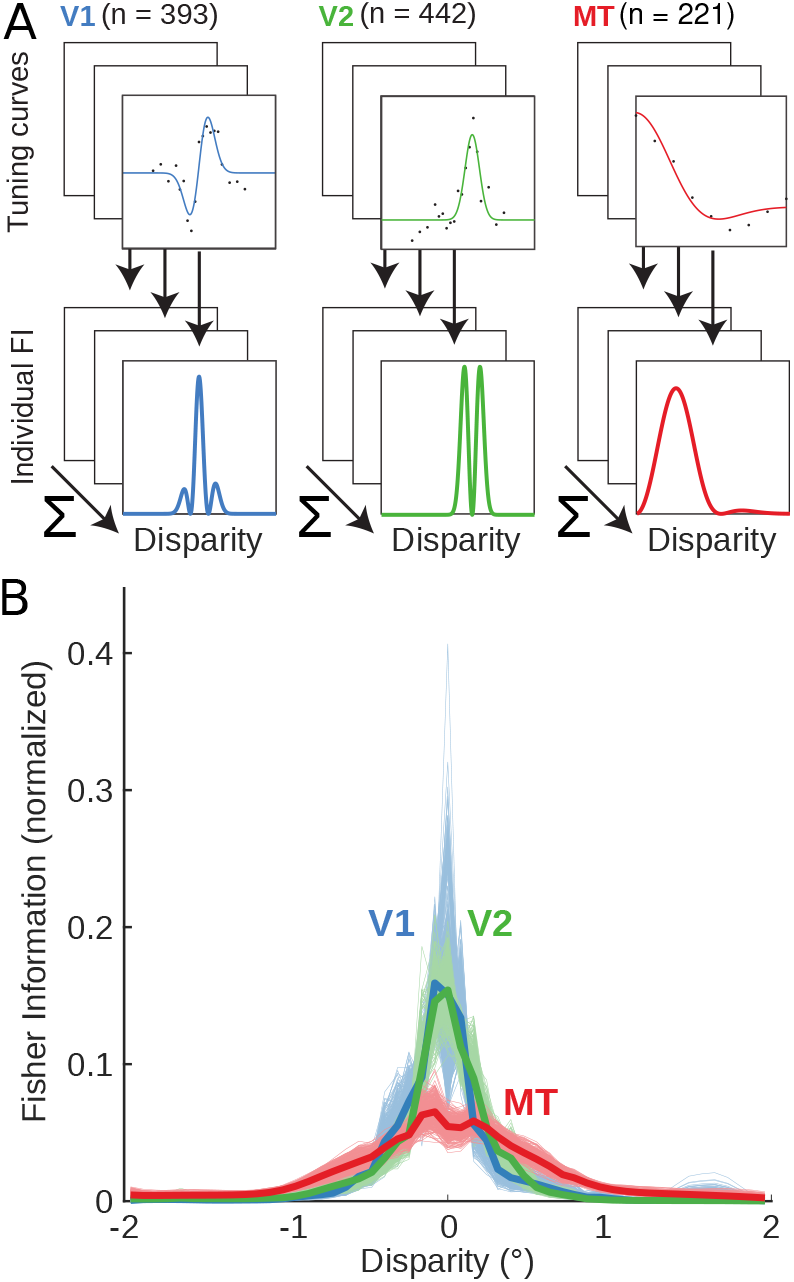
**A.** We compiled a data set of 1056 disparity tuning curves from brain areas V1, V2, and MT of the macaque monkey. The mean responses of each neuron as a function of disparity were fit with a continuous function and the individual neuron *FI* associated with each tuning curve was calculated from these fits. **B.** The population *FI* (see Fig. 1C) is shown for each brain area (thick lines). Thin lines represent the population *FI* computed from 500 bootstrapped samples from each brain area.

For a single neuron, the *FI* is high when the tuning curve is steep and the spike rate (and Poisson noise) is low, and the *FI* is low when the tuning curve is flat and the spike rate is high (for example, note the alignment between the tuning curve flanks and the *FI* peaks in Fig. 2A). The total *FI* of each population was calculated as a sum across neurons, based on the assumption that each neuron responds to stimuli independently (Fig. 2B). Qualitatively, we see that the population *FI* is most kurtotic in V1/V2 and least kurtotic in MT, consistent with the hypothesis that the information-maximizing model is a better description of the early visual representation (V1 and V2) and the discrimination-maximizing model is a better description of the downstream representation (MT).

We next directly compared these empirical *FI* distributions to the two probability distributions of binocular disparities in the natural environment. Binocular disparities are distributed non-uniformly across the visual field during natural behavior (12), so we started by resampling the natural disparity distributions based on the specific retinotopic locations of neuronal receptive fields in each population using kernel-smoothed density estimates (see Fig. S1). The resulting disparity distributions were all similar in shape (Fig. S2A&B), despite the minor differences in sampling density between the different brain regions.

We calculated the power law that, when applied to the corresponding disparity distributions, resulted in the best fit to the measured *FI* of each neuronal population. We used bootstrapping of the populations to estimate variability of the best-fit power law exponent. Consistent with our working hypothesis, we observed a systematic decrease in the distribution of exponents for the best fit power law from the lower-level areas V1 and V2 to mid-level area MT for both natural tasks (Fig. 3A&C). Note that in both plots, the distributions for V1 and V2 are largely overlapping. The results of these fits suggest that V1 and V2 are closer to populations optimized to preserve information about binocular disparity (particularly for the food preparation task, for which the best fit exponents are 1.64 and 1.57, respectively). The best fit exponent for MT was consistently lower for both tasks, and therefore less consistent with information maximization and more consistent with reducing error magnitude. Of course, the differences between V1/V2 and MT are not as extreme as the infomax and discrimax examples given in Fig 1, but the relative shift is robust and persistent across both example disparity distributions.

**Fig. 3.**
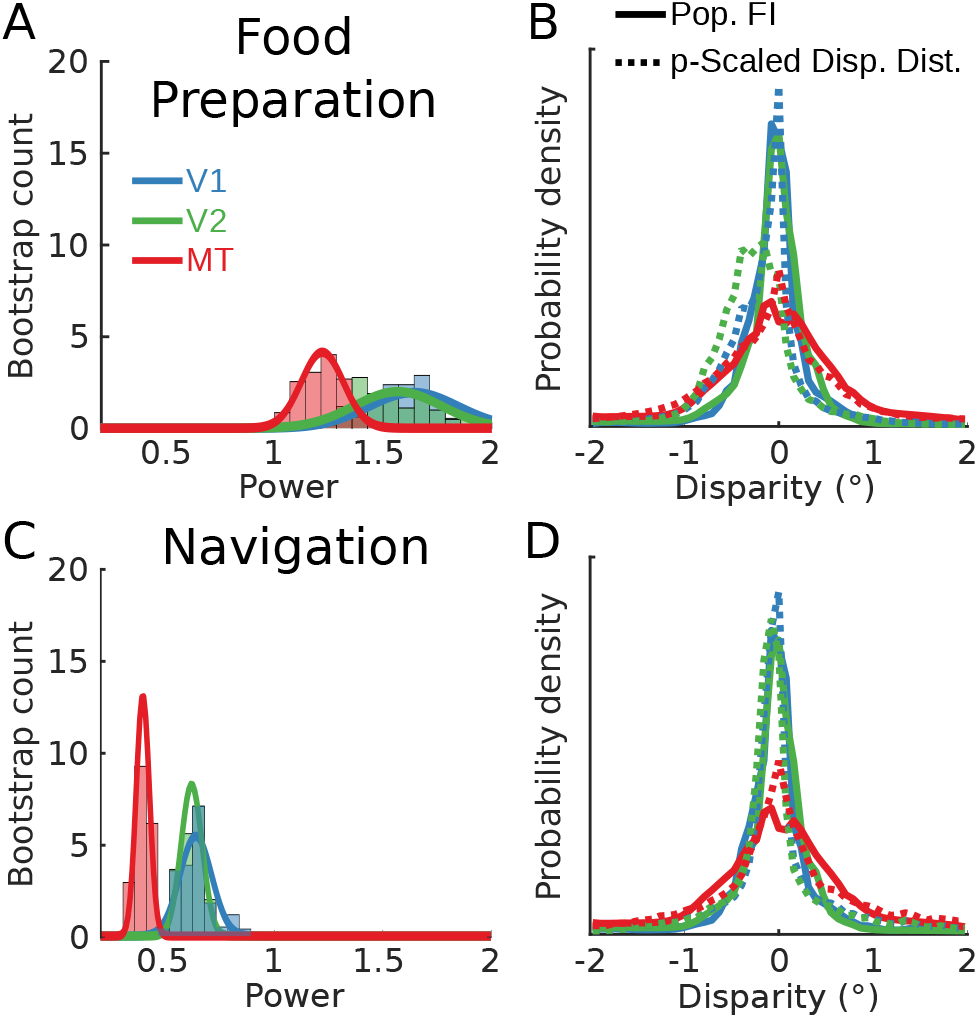
**A.** The distribution of best-fit power law exponents linking population *FI* to the disparity probability densities is plotted for the food preparation data set (using the kernel-smoothed probability densities to guide sampling). Histogram bars indicate the 500 bootstrapped samples for each brain area and solid lines indicate the Gaussian distribution fits to each set of samples. Note that the V1 and V2 distributions are highly overlapping. **B.** Population *FI* is plotted for each area (solid lines) along with the disparity distributions scaled by their respective best fit power laws (dotted lines; see Fig. S2)). The power law exponents are as follows: V1: 1.51, V2: 1.69, MT:1.22. **C & D.** As in **A & B.**, but for disparity statistics collected during the navigation behavioral task. Power law exponents: V1: 0.61, V2: 0.62, MT: 0.41.

However, we observed one notable inconsistency with this interpretation, with respect to V2 and the food preparation task. Fig. 3B&D show the matches between the population *FI* (solid lines) and the binocular disparity distributions scaled by the single best fit exponents (dashed lines) from Fig. 3A&C. According to our predictions, for each brain area and each task these pairs of distributions should closely overlap. The distributions match closely in all cases except for the V2 population and the food preparation task (green lines in Fig. 3B). The power-law scaled binocular disparity distribution in this case is quite biased towards near (negative) disparities, and this bias is not present in the population *FI*. The reason for the near disparity bias is clear from the distribution of receptive field locations (Fig. S1): the V2 receptive fields are exclusively concentrated in the lower visual field, and the disparity statistics in the lower visual field are strongly biased towards near disparities during food preparation (12). For the navigation task, the binocular disparity statistics are less biased. At present, we do not understand how the visual system might flexibly incorporate different biases in stimulus statistics when they differ notably across different tasks. However, we speculate that the notable lower visual field near bias in this food preparation task (in which participants made a sandwich while sitting at a table) may not be strongly reflected in the visual experience of macaque monkeys.

To quantify the exponent differences further, we fit the bootstrapped power law values for each neuronal population with a Gaussian distribution and measured the effect sizes between populations. The effect sizes, measured as Cohen’s D between pairs of populations, were large between the earlier areas and MT (food preparation: V1 vs. MT = 2.6, V2 vs. MT = 2.2; navigation: V1 vs. MT = 4.3, V2 vs. MT = 5.5). As expected from the similarity in their *FI* distributions, the effect sizes were small between V1 and V2 (food preparation: 0.36, navigation: V1 vs. V2 = 0.31). Thus, the data support the notion that the MT *FI* reflects a different optimization but that V1 and V2 may contain similar information-driven codes.

### These differences correspond to a broad set of changes in individual tuning curve characteristics

### from V1/V2 to MT

We next asked which aspects of the neural responses to disparity could account for the differences in the *FI* distributions between V1/V2 and MT. To answer this question, we first took a parametric approach and leveraged the Gabor fits to each of the tuning curves (Fig. 4A). For each of the cortical populations, we examined the distributions of each of the six best-fit Gabor parameters (Fig. 4B). In sum, we find that MT neurons generally have higher response offsets, broader envelopes, and lower disparity frequencies than either V1 or V2. The MT population also has a broader distribution of best fit envelope means and phases of the cosine component than the earlier cortical areas. Full results of the statistical comparisons across populations are provided in Tables S1 & S2. This analysis expands on a previous comparison between V1 and MT (22) by including responses from V2 and a larger number of V1 neurons. One possible explanation for these differences is that they may reflect differences in the retinoptic locations of the receptive fields across the samples from each brain areas. Our subsampling from the initial larger data set resulted a good match between the brain areas in terms of eccentricity and vertical position within the visual field, although the MT data set is more concentrated in the left visual field and the V1/V2 data sets are more concentrated in the right visual field (see Fig. S1). Since there is no reason to hypothesize that disparity tuning should differ in terms of left or right visual field, these tuning differences most likely reflect differences in the underlying neural representation.

**Fig. 4.**
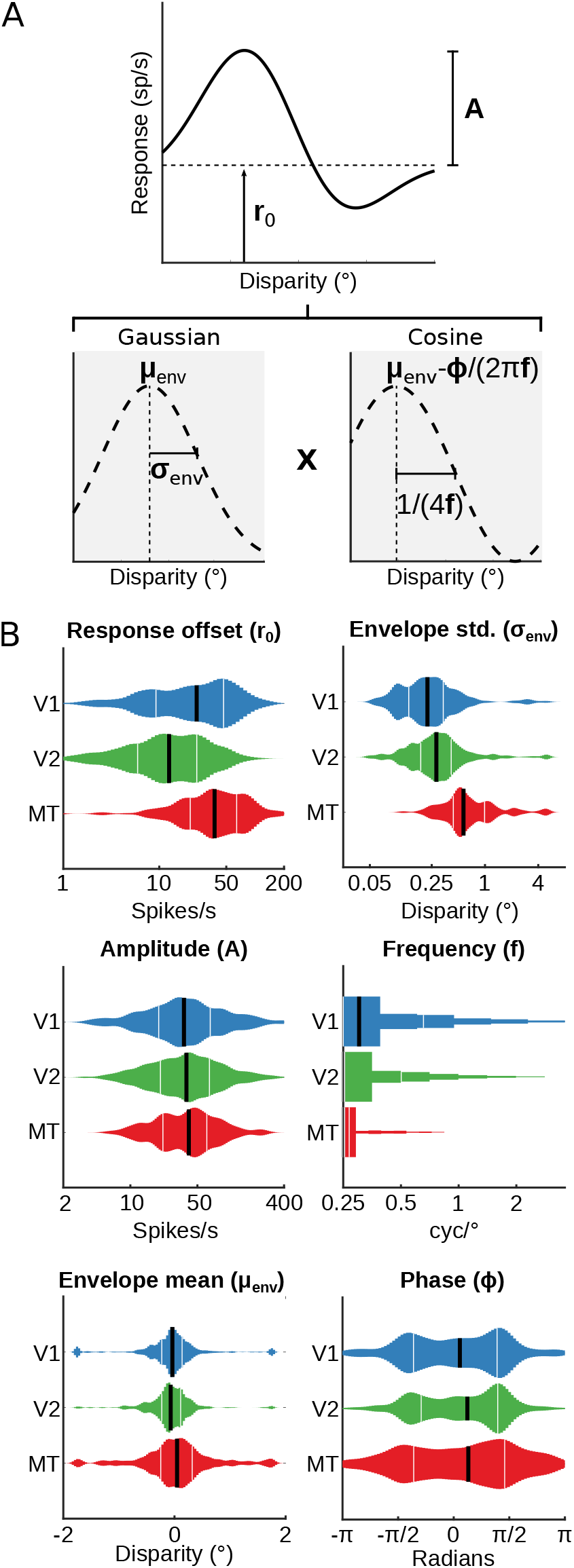
We fit all the neuronal data with a modified 6 parameter Gabor function and plotted the distribution of best-fitting parameters for each cortical area from which we have data. **A.** Decomposing the Gabor function into Gaussian (left plot) and cosine (right plot) components clarifies what each parameter contributes to the shape of the resulting tuning curve. The vertical dashed line in the Gaussian plot marks the center of the Gaussian envelope while the cosine plot indicates the 0 phase position. The parameters are defined as follows: *A*: Amplitude, *r*_0_: vertical response offset, *μ_env_*: Gaussian envelope mean, *σ_env_*: Gaussian envelope standard deviation, *f*: cosine frequency, *ϕ*: cosine phase. The cosine *f* parameter is shown in the bottom right panel as defining one quarter of a period. **B.** Distributions of best fitting Gabor parameters for each of the 3 cortical areas. Thin white bars indicate the 25th and 75th quartile and the thick black bar indicates the median.

### An increase in neurons preferring larger disparities is a key factor in the observed coding transformation

There are clear differences in the distribution of best fit Gabor parameters and the preferred disparities between the earlier cortical regions and MT. However, the complex and overlapping effects that these parameters have on the tuning curve shape make it hard to interpret how each parameter contributes to the transformation of the population *FI* distributions between the regions. Therefore, we performed a resampling analysis to see if the changes in any one parameter in particular could account for the difference in the *FI* distributions between V1 and MT. Since the *FI* distributions from V1 and V2 were similar, we did not repeat this analysis for the difference between V2 and MT. The overall approach is outlined in Fig. 5A: for one tuning curve parameter at a time (illustrated just for frequency), we replaced the set of true V1 values with a new set of values obtained by randomly sampling from the distribution of MT fits. We then rebuilt each cell’s tuning curve with their new parameter and used this to calculate a new *FI* distribution for each cell. Lastly, we summed the individual cell *FI* distributions and normalized by the area under the curve to compare the overall shapes of the resampled *FI* distribution for V1 and the true *FI* distribution for MT. This process was repeated a total of 500 times for this Gabor parameter and then another 500 times for each of the Gabor parameters individually to assess the variability in the resulting resampled V1 population *FI* distributions. Fig. 5B shows the resulting V1 population *FI* distributions (black) for each of the parameters alongside the true MT population *FI* distribution (red). Of the six parameters, replacing the V1 envelope mean parameter with those from MT qualitatively results in the closest match with the lowest variability.

**Fig. 5.**
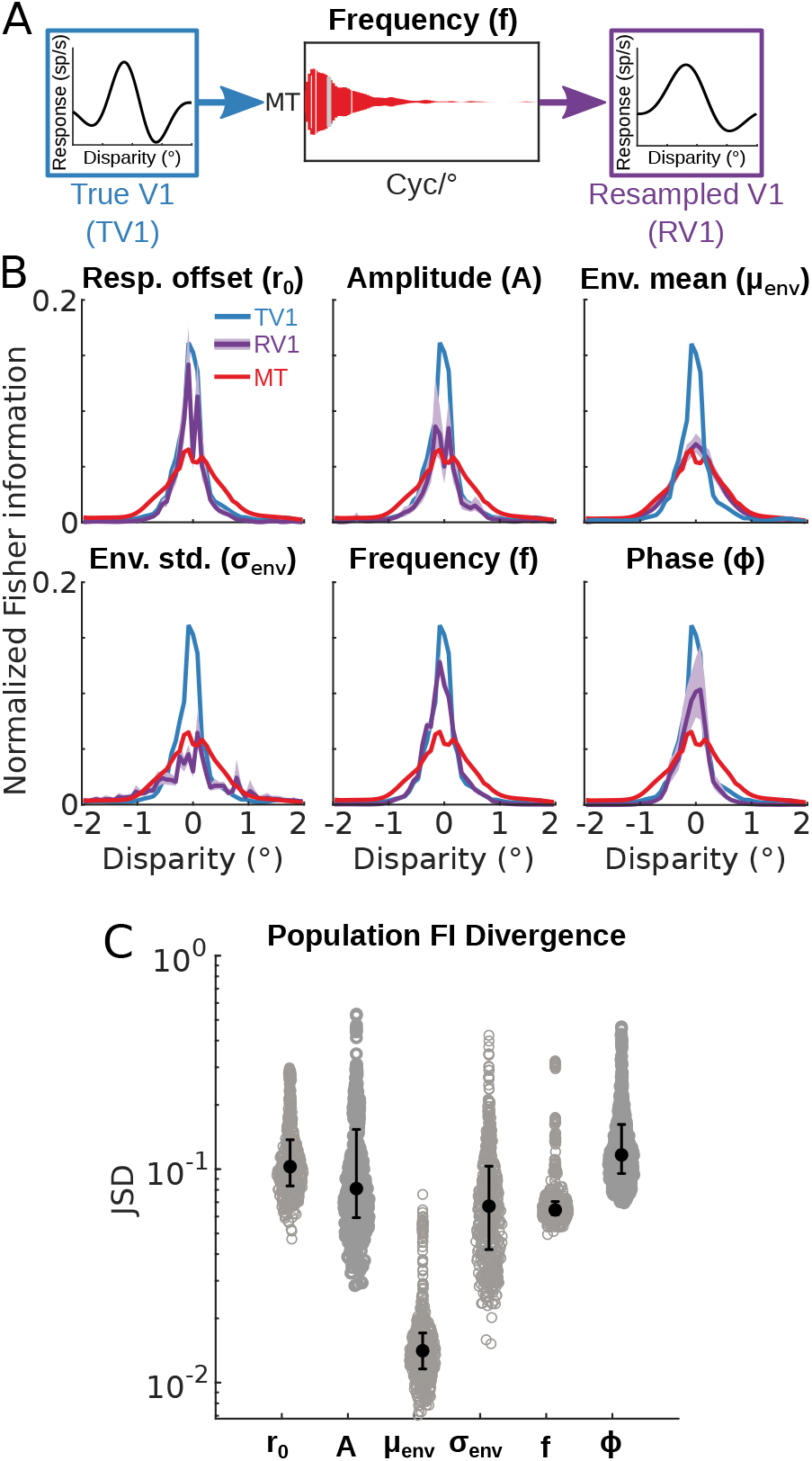
**A.** We investigated which of the 6 Gabor parameters best explained the difference in the population *FI* between V1 and MT by replacing (one parameter at a time) from V1 with randomly sampled values pulled from the distribution of best-fitting parameters from the MT population. We repeated this process to create 500 new bootstrapped V1 populations for each of the 6 parameters. **B.** The mean and interquartile range of the population *FI* distributions from the 6 resampled V1 populations (purple) are plotted against the true MT population *FI* (red, same in each of the 6 subplots). The original true V1 *FI* distribution before resampling is shown as blue lines for reference (same in each of the 6 subplots). **C.** Jensen-Shannon Divergence (JSD) between the resampled V1 population *FI* distributions and the true MT population *FI* distribution. Lower values reflect a closer match between the two distributions. (Gray circles) JSD between the population *FI* for each bootstrapped populations (i.e. RV1) and the true MT population *FI*. (Black circles, errorbars) Median JSD and interquartile range across all bootstraps.

To examine the significance of these matches, we calculated the Jensen-Shannon divergence (JSD) between the *FI* of each of the resampled V1 populations and the true MT population *FI* distribution (Fig. 5C). This information theoretic measure reflects the dissimilarity between two distributions. We first tested whether there were significant differences between the sets of JSDs for each parameter using a Kruskal-Wallis test and confirmed there were differences in dissimilarity between the parameters (*χ*^2^ = 1.65E3; *df*=5; *p*<2.22E-16). While there were significant differences between each of the sets of resampled populations (see Table S3), the *μ_env_* parameter stood out in that it’s associated JSD values were substantially lower than all other parameters (note the large z-scores) – that is, it was the least divergent from (most similar to) the empirical MT distribution. We performed two control analyses to examine the generality of these results (both analyses are described in the Methods and reported in Fig. S4 and Table S5 and S4).

From this pattern of results, we can glean that much of the difference in how disparity information is represented between earlier and later cortical areas can be attributed the presence of a greater number of cells that are selective for larger disparities in MT.

## Discussion

Here we have taken a theoretical prediction about population-level neuronal information along the sensory processing hierarchy and put it to an empirical test. The results are consistent with the prediction that sensory transformations can be understood as the result of a constrained optimization, in which the goal changes sensibly from early to late sensory regions.

While we focused on the power law relationship between stimulus probability, encoding goals, and population *FI*, the optimal population *FI* can be influenced by other factors as well. For example, the optimal power law is affected by the resource constraints (21). We assumed that visual brain regions in the same species are subject to the same constraints. Adaptation studies that dynamically shift stimulus statistics without affecting constraints may be able to determine whether or not this assumption holds (23). Similarly, system noise influences the form of *FI*, both for individual neurons and the population as a whole (see (24)). A noise analysis of our data suggests that, at least in the current data set, spike count properties were similar across the examined brain areas (see Fig S3). At the population level, our *FI* calculation assumes that the neurons in our populations are independent, but we cannot confirm the validity of this assumption via measurements of spike count correlations between neurons since our data consists primarily of single unit recordings. Response correlations reduce population *FI* in many cases, but it isn’t clear whether the magnitude of spike count correlations vary between the cortical areas analyzed here. A systematic characterization of neuronal noise properties and noise correlations along the sensory processing hierarchy may ultimately reveal that sensory transformations are subject to differing noise properties as well (25, 26).

Here, we used binocular integration as a model system for sensory transformations more generally. But our results also shed light on specific open questions in the neural underpinning of binocular integration and binocular disparity processing. Previous work has noted that the shape of disparity tuning curves appears to change systematically across brain regions, but the reason for these changes is unknown (22, 27, 28). For example, previous work suggested that MT tuning curves tend to have odd symmetry and broader tunings, whereas in V1, the best fit tuning curves are more even symmetric with a narrower range of preferred disparities (22). Here, we did not see evidence for a difference in even/odd symmetry, but did observe a multifaceted set of differences in the distributions of tuning curve shapes. Some of the difference in tuning shape (e.g., width) may be a side effect of pooling neurons with different orientation preferences to generate direction selectivity for patterns (29). However, a priori it is not necessarily the case that tuning curves get broader along the sensory processing hierarchy. For example, the tuning curve characteristics in V2 were quite similar to V1. Our information theoretic analysis suggests these differences in tuning curve shape may also have direct utility: they shift the position of a neuron’s peak *FI* to larger disparities while maintaining a population peak near zero. Over the entire population, this effectively makes the population *FI* distribution more broad, thus improving disparity discrimination at higher disparity pedestals.

Of note, we do not claim that a shift from “infomax” to “discrimax” representations is the sole difference in how V1, V2, and MT represent visual information. For example, it is well-established that MT encodes binocular disparity in a way more correlated with perception than V1 does: V1 neurons invert their disparity tunings with anticorrelated stereoimages (7) and are affected by vergence and absolute disparity (30, 31), whereas V2 (32, 33) and MT (34) at least partially discard false stereo matches and encode relative disparity. It is also important to consider other sensory variables represented in these areas. A recent study used an information theoretic framework to examine the link between the encoding of speed in MT and perceptual biases in speed estimation (35). They found that the population *FI* for stimulus speed in MT can be related to speed perception via a *p*^2^ power law. Our estimates of the power law in MT for binocular disparity diverge from this idealized “infomax” representation of speed. However, it is not necessary that every sensory variable encoded within a population is represented at the same level–it is entirely possible that the same brain region could contain an information-maximizing representation of one sensory variable and a discrimination-maximizing representation of another. The information-theoretic framework provides an additional window into how neural representations build and interact along sensory processing streams that can complement other assessments of neural function.

Lastly, our work also highlights the fact that the statistics of sensory input can be task dependent. The binocular disparity statistics were quite different for the two tasks that we considered. This task-dependence can pose a problem for assessing the encoding optimality of neural populations on the basis of task-free natural stimulus statistics derived from generic data sets, for example, of natural images and sounds (3, 4). Here, we show that generalizations that are robust across tasks can be made by focusing on relative differences between brain regions.

With the information theoretic analyses like those presented here, we gain a more principled understanding of the link between the hierarchy of cortical areas carrying sensory information and the complexity of behaviors that rely on that information. New computational frameworks that can be applied to dynamic populations of neurons, trial-by-trial variations, and spike-count correlations between neurons will contribute to the next step in characterizing hierarchical transformations of sensory signals.

## Methods

### Definition of Horizontal Binocular Disparity

We define horizontal binocular disparity (*d*) as follows:

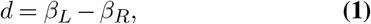

where *β_L_* and *β_R_* denote the horizontal angular eccentricity of an image projected to the left and right retinas, relative to the fovea. We represent eccentricity and binocular disparity in units of visual degrees, with negative disparity values indicating points that are closer in depth than the fixation point and positive values indicating points that are farther in depth than the fixation point.

### Natural Scene Statistics of Binocular Disparity

Natural statistics of binocular disparity were re-analyzed from a previously collected data set (12). In brief, three adult human subjects performed either indoor navigation or food preparation tasks while wearing a custom-designed headset consisting of a pair of stereocameras and eye trackers. Still images were sampled from the stereocamera video footage, transformed into head-centered 3D scene geometry, and finally converted into binocular disparity maps in retina-centered coordinates using the eye-tracking data. Analyses were limited to data within a 10° radius of the point of fixation.

While the original natural binocular disparity statistics measurements were evenly sampled from a 10° radius of fixation, the neural data sets contained neurons with receptive fields that were restricted largely to one hemifield. Given the differences in binocular disparity statistics between the upper and lower hemifields and the expected increase in prevalence of larger disparities with increasing eccentricity (12), these biases on receptive field location likely influence the probability distribution of binocular disparities that each neuronal population encodes. Thus, we resampled binocular disparities from the maps based on the receptive field centers of each neuron. To do so, we calculated 2D kernel-smoothed probability density distributions (see Fig. S1) for each of the cortical areas using a 2D Gaussian kernel and used these distributions to subsample from the original binocular disparity distribution. To ensure the reliability of this subsampling procedure, we used a bootstrapping procedure to estimate the variability of the resulting distribution (errorbars are smaller than the line width in Fig. 3A&D). In short, we sampled with replacement from each of the binocular disparity data sets (navigation and food preparation) to generate 100 bootstapped samples of equal size to the empirical distribution, calculated the disparity probability distribution for each bootstrap, and recovered the 95% CIs from the cumulative error distributions.

### Neural Recordings

Neural recordings from areas V1, V2, and MT were re-analyzed from a combination of multiple previous studies. All recordings come from awake fixating macaque monkeys and reflect measured action potentials of isolated neurons. The tuning curves measured in V1 and V2 come from multiple studies using highly similar methods measured over the course of a decade in the same laboratory (36–46). The MT tuning curves were obtained in a single study with slightly different experimental methods (22).

In all cases, the stimuli used to measure response rate (spikes per second) as a function of binocular disparity were random dot stereograms (RDS) with each dot subtending approximately 0.1°. The methods for the stimuli and data collection methods for the V1 and V2 data are described in (37, 40). Briefly, responses from V1 and V2 were collected using RDS stimuli with no coherent motion, presented on a Wheatstone mirror haploscope for 400-500ms at a display refresh rates ranging from 72-100Hz. Responses from MT were collected using RDS stimuli with 100% coherent motion tailored to each cell’s preferred direction, speed, and size (22). For each of the MT recording sessions, stereoscopic stimuli were presented for 1500ms on a single monitor with liquid crystal shutter glasses at a refresh rate of 50Hz for each eye (crosstalk was measured to be <3%). The stimuli in the MT recording sessions were also presented against a non-overlapping background of stationary dots at zero disparity to anchor vergence (this vergence lock was located outside of the neuronal receptive fields in all cases).

Recordings from each area were likewise similar. In brief, after the receptive field of each neuron was localized, the stimulus was adjusted to the optimal size (and velocity for MT) and the responses to different magnitudes and directions of binocular disparity were recorded. For the MT recordings, the majority of tuning curves were mapped using stimuli with disparities that ranged from −1.6° to 1.6° in steps of 0.4°, however for a few neurons larger disparities were used. For the V1 and V2 recordings, the stimulus disparity ranges were variable across neurons, with steps ranging from 0.029° to 1.2°. Responses from MT and V2 neurons were collected solely with single electrodes, while responses from V1 were collected mostly from single electrodes and a few from multicontact probes (45). The receptive fields of neurons in V1 and V2 were largely restricted to the lower visual field due to cortical topography and the positioning of the implanted recording cylinders. In the MT data set, most recording cylinders were placed above the right hemisphere, so the receptive fields are largely limited to the left visual field. Horizontal and vertical eccentricity of the stimuli and receptive field positions were calculated by taking the arctangent of the distance on the screen from the fixation point and the viewing distance.

There were slight differences in experimental protocols between the laboratories (i.e., shutter glasses vs. mirror haploscope, stimulus presentation time, coherent motion vs. incoherent motion), but these differences are unlikely to cause the differences in the *FI* distributions estimated from the data. For example, Palanca and DeAngelis (2003) compared disparity tuning in MT for moving and stationary dots, and the disparity tuning curves were very similar (although responses were generally weaker for stationary dots) (47). Indeed, motion and disparity are independently encoded in MT (48), so it is unlikely that the difference in motion energy between the stimuli used by the two laboratories would produce consistent biases in disparity tuning. Most neurons in V1 and MT also do not show a dependence of disparity selectivity on interocular delay, instead showing an inverse relationship between response gain and interocular delay; those that do show disparity-delay inseparability do not exhibit large tuning shifts over the display intervals used in either of the data sets (40, 49).

### Tuning Curve Analysis

A subset of the neural recordings described in the previous section were selected for analysis according to set a of inclusion criteria. First, we only included neurons with an average of ≥ 3 repeats per stimulus disparity. We then selected neurons for which a significant amount of the trial-by-trial variance in responses was explained by the disparity of the stimulus (one-way ANOVA at a significance level of p < 0.01). For the V1 and V2 recordings, the range of stimulus disparities presented was variable, so to ensure sufficient data to obtain a reliable fit to the tuning curves we only analyzed neurons with a range of at least 1° between the nearest and the farthest disparity tested. These criteria resulted in 690, 531, and 444 neurons from V1, V2, and MT, respectively.

In each study, stimulus disparity was recorded in screen coordinates rather than retinal coordinates. For example, a point with the same horizontal coordinate on screen for both eyes was coded as having zero disparity. However, in retinal coordinates, the locus of points with zero disparity is a circle that contains the fixation point and the optical center of the two eyes. Therefore, a point with the same horizontal coordinates on a planar screen will have an uncrossed (far) retinal disparity. In order to match up the neural recordings to the retinal disparities measured in the scene statistics analysis, we therefore applied a correction factor. In brief, the retinal eccentricity of each on screen stimulus in the left and right eye was determined based on the screen distance, the horizontal screen coordinates in the left and right eye, and an assumed interocular separation of 30mm. These retinal eccentricities were used to calculate the angular disparity on the retinas, which were were used in the subsequent analyses.

For each of these neurons, we then fit a continuous tuning curve to the mean responses as a function of the stimulus retinal disparity *h*(*d*). Tuning curves were parameterized as a 6-parameter Gabor function:

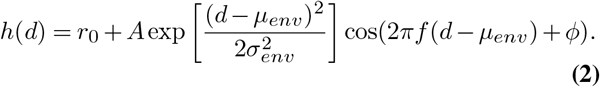

Best-fitting parameters for each neuron were determined using constrained non-linear optimization in MATLAB (Mathworks, Inc.), minimizing the mean squared error (fmincon). To prevent fits that deviated substantially from the observed range of spike rates, data were up-sampled by a factor of 2 using linear interpolation prior to fitting. Bounds for parameters were as follows: 0 < *r*_0_ < 500, 0 < *A* < 500, −1.75 < *μ_env_* < 1.75, 0 < *σ_env_* < 5, 0 < *f* < 4.5, –2*π* < *ϕ* < 2*π*. Fits that resulted in a minimum spike rate of less than 0.05 spikes per second or with a frequency (*f*) of less than 0.25 were strongly penalized (by multiplying the current error by 1*E*7), to avoid instability in the calculation of Fisher Information and the interpretation of the fitted parameters, respectively. The optimization was initialized at 200 randomly selected starting points and optimized according to an interior point algorithm. The parameters with the lowest error across all initializations were taken as the final fit. A subset of neurons (less than 30) were identified with poor fits on manual inspection, so fitting routines were re-run for these neurons with minor adjustments to the parameter ranges. The *R*^2^ across all fits was greater than 0.3, and the median was above 0.8 for all three areas.

We further subsampled the neuronal populations to ensure that each included cell was well-fit with a Gabor tuning function (we select only neurons with *R*^2^ ≥ 0.75), had RF centers within the defined region of the disparity image set (i.e. ≤ 10° eccentricity), and had RF centers within the same general subregion of the visual field. To enforce the final criterion, we determined the largest vertical and horizontal components of the RF centers from the V1/V2 data sets to define a rectangular bounding box and then selected only MT cells with RF centers within this region. This was enforced because the MT data set contained a larger number of cells with RF centers in the upper visual field, which could potentially bias the disparity preferences of the sample (12). The final cell counts from each sample were 393, 442, and 221 from V1, V2, and MT, respectively.

To investigate differences between the resulting parameter distributions, we first performed a Kruskal-Wallis omnibus test to ask if there were any significant differences in median values between the three cortical regions. Since we were primarily interested in differences in distribution breadth, we first took the absolute value of the signed parameters (*μ_env_* and *ϕ*) to test only difference in the median magnitudes of these values. If the omnibus test revealed a significant difference between the distributions, we then followed up with a set of Wilcoxon rank sum tests between each of the pairs of cortical areas. The results of these tests are presented in Tables S1 and S2.

### Fisher Information

Based on the simplifying assumption that the spike rates for each neuron are Poisson-distributed, we calculated the Fisher Information (*FI_n_*) of each neuron as described by (50):

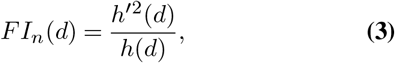

where *h′*(*d*) denotes the first derivative of the tuning curve. Negative *FI_n_* values, resulting from fits with small negative spike rates, were set to zero. Assuming that each neuron’s spike rate is independent, the *FI* in the population level is then:

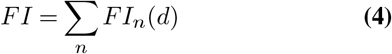

Thus, we summed together the *FI_n_* across the neurons in each neuronal population.

To examine the variance of the population *FI*, we repeated this analysis by bootstrapping samples of 200 neurons (sampled with replacement) 500 times for each region. The sample size of 200 was selected so that we could match sample sizes across all three populations.

To ensure that our neuronal data sets contained disparity responses that were approximately Poisson and did not substantially vary between areas, we plotted the relationship between mean spike count and spike count variability for each of the cell populations (Fig. S3, top). Note that each point is a unique presented disparity condition and each cell contributes multiple points. Only conditions with ≥ 5 repeated stimulus presentation were included. We then fit a simple power law function to each of the distributions:

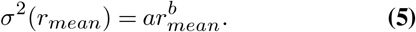

Poisson-distributed responses should be best fit with both *a*, or *slope*, and *b*, or *power law*, equal to 1. While the best fit slope for each area is slightly greater than 1, the best fit power laws did not substantially differ from 1 (Fig. S3, bottom). Since the populations also did not greatly differ in their best fit values, our assumption of Poisson spiking statistics for the calculation of population *FI* seems well-justified.

### Gabor Parameter Resampling Between Cortical Areas

To identify what aspect of disparity tuning could best explain the differences in the *FI* distributions between populations, we conducted a resampling approach whereby we replaced the best-fitting Gabor parameters individually for each parameter (*r*_0_, *A, μ_env_, σ_env_, f, ϕ*) from the set of cells in V1 with parameters sampled from the distribution of fits from the set of cells in MT (diagrammed in Fig. 5A). Since the *FI* distributions from V1 and V2 did not significantly differ, we restricted the analysis to V1 and MT. For each parameter, we first discarded the best-fit set from the V1 population. We then randomly sampled from a kernel-smoothed probability density derived from the set of best-fits to MT neurons and assigned new values to the parameter of interest for the V1 population. As done previously, we then calculated the single cell *FI* distributions given this new set of tuning curves and summed their values to get the population FI. We then repeated this process 500 times to obtain confidence intervals on the population Fisher information under the reparameterization. This process was then repeated for each of the Gabor parameters individually, with the median *FI* distributions and interquartile range for the resampled V1 populations shown in Fig. 5B. To quantitatively determine which resampled parameter produced a V1 *FI* distribution that was closest to the empirical MT distribution, we computed the Jensen-Shannon divergence (JSD) between each of the bootstrapped V1 *FI* distributions and the empirical MT distribution.

We performed two control analyses to examine the generality of these results. First, we wanted to ensure that our focus on comparing population *FI shape* similarity instead of comparing the similarity in *area under the curve* (AUC) of the population *FIs* did not lead us to a false conclusion about the influence of each parameter. Therefore, we repeated our analysis without normalizing each bootstrap by the AUC and instead normalized solely by the number of cells in the bootstrapped sample. Overall, the results again show a large *FI* similarity between the resampled *μ_env_* parameter populations and the true MT *FI* distribution (Fig. S4A & black/gray data points in Fig. S4B). Note, however, that this metric is not very sensitive to differences in the distributions at larger disparity values where probabilities are low (resulting in deceptively close AUC values between the true MT population *FI* and the resampled *f* parameter population *FI* as well). To ensure that the results of our resampling approach were due to resampling from the MT parameter distributions in particular and not due to simply shuffling the best fit V1 parameters, we also repeated our AUC and JSD analyses by resampling from the V1 parameter distribution instead of MT (i.e., we shuffled the parameters between V1 cells with replacement). We plot the results of these analyses alongside the MT parameter sampling results for the AUC and JSD metrics in Fig. S4B & D, respectively (orange data points). We examined significant differences between the populations sampled from the V1 and from the MT distributions by performing a Wilcoxon rank-sum test for each Gabor parameter. The results of these tests are shown in Table S4 (AUC) and Table S5 (JSD) and significant differences are indicated with asterisks in Fig. S4B & D. Comparing the results between the bootstrapped V1 populations that sampled parameter values from MT and those that had their parameters shuffled between V1 cells, we find that sampling from MT significantly minimized the divergence in population *FI* between MT and V1 for the *μ_env_*, consistent with the interpretation from our main analysis. More specifically, it shows that the *specific* changes in the distribution of preferred disparities in MT greatly contributes to the change in *FI* distribution between V1 and MT.

### Comparison Between Fisher Information and Disparity Statistics

We used a grid search to determine the power law that minimized the difference between each population *FI* and the sampled binocular disparity probability distributions. Power laws were applied to the disparity distributions and then differences were calculated as the mean absolute error between the two distributions sampled at 51 evenly spaced binocular disparities between –2° and 2°. We repeated this minimization for each of the 500 bootstrapped samples from each neuronal population and fit the resulting distributions with a Gaussian distribution using maximum likelihood estimation, which also allowed us to characterize the 95% confidence interval of the average power law for each neuronal population.

## Code and Data Availability

The MATLAB code used to pre-process the neuronal data, perform the neuronal analyses, and collect the image statistics is available at https://github.com/tsmanning/DisparityInfoProject. The neuronal data and BORIS data sets are available at [Redacted until final publication].

## Acknowledgements

We thank the National Eye Institute (grants F32 EY032321, T32 EY007043, and R01 EY022443) for support.

