## Supplementary Information for "Transformations of sensory information in the brain reflect a changing definition of optimality"

Tyler S. Manning

Emma Alexander

Bruce G. Cumming

Gregory C. DeAngelis

Xin Huang

Emily A. Cooper

**Corresponding Author:** Tyler S. Manning

**This PDF file includes:**

Figures S1 to S4

Tables S1 to S5

### Kernel-smoothed density distributions

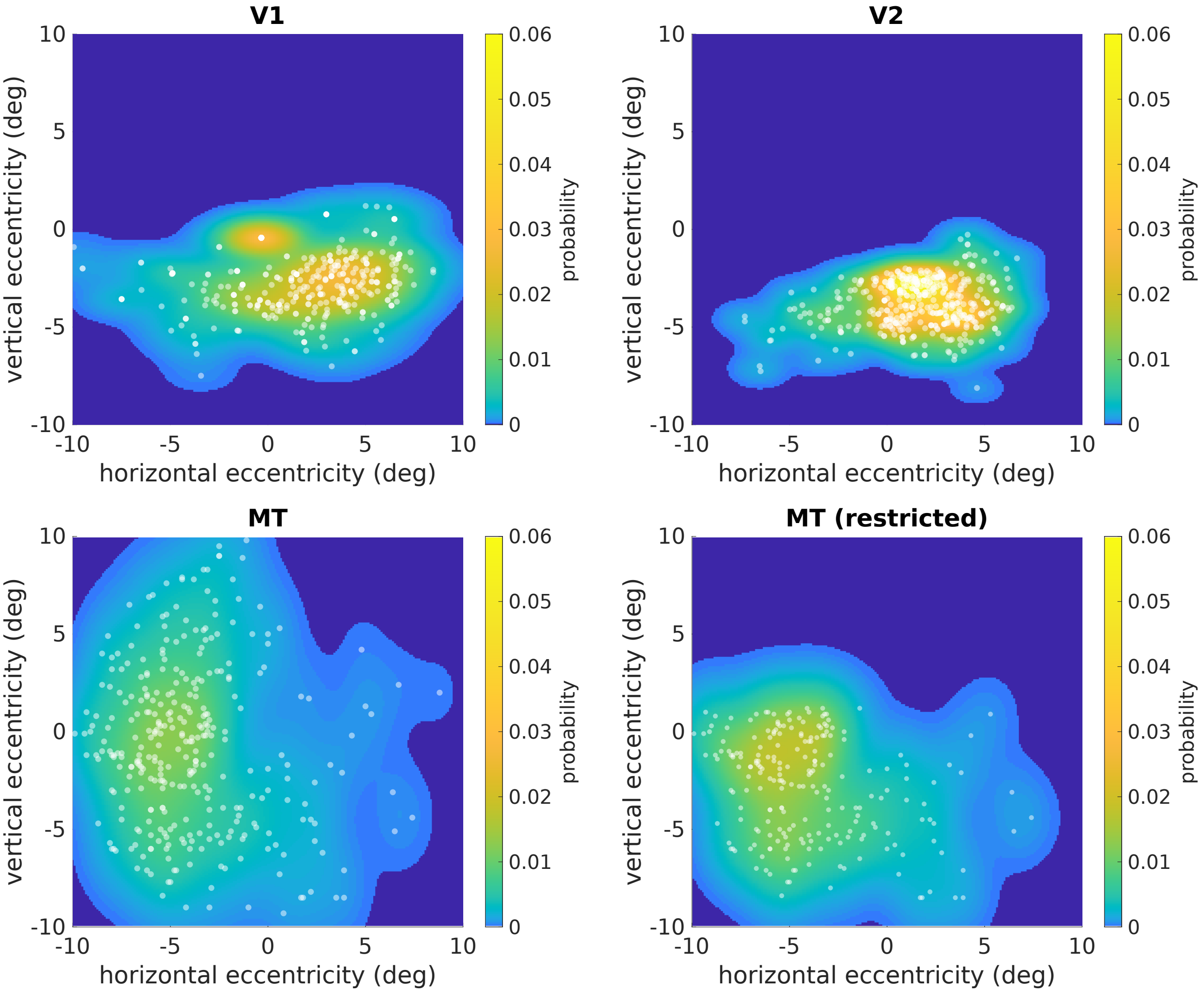


**Fig. S1.** The distribution of receptive field retinotopic locations varied across brain areas. The panels labelled V1, V2, and MT show the locations of the centers of all receptive fields (white dots) in each brain area as a function of horizontal and vertical eccentricity. Kernel-smoothed probability densities (KSDs) are shown as heat maps. The final panel, labelled MT (restricted) shows the receptive field centers and KSD for the subset of MT neurons used in the analysis. This MT subset was restricted to neurons with receptive field centers within the bounds of the maximal extents of the V1/V2 receptive fields.

### Disparity probability densities sampled with KSDs

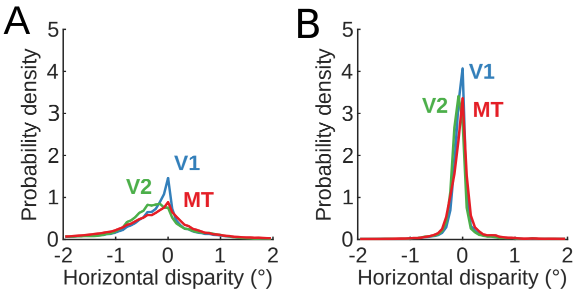


**Fig. S2.** Disparity probability distributions differ across the visual field, so we tailored a set of three probability distributions to the kernel-smoothed density of the receptive field centers in each neuronal population (see Fig. S1). **A.** The disparity probability densities collected from the food preparation task weighted based on the spatial sampling of the V1 (blue), V2 (green), and MT (red) data sets (see Fig. S1). The mean and 95% confidence intervals (CI) of the disparity probability distributions across 100 bootstrapped samples are shown, but the CI error bars are too small to be visible. **B.** As in **A.**, but for the navigation task

### Spike count statistics between cortical areas

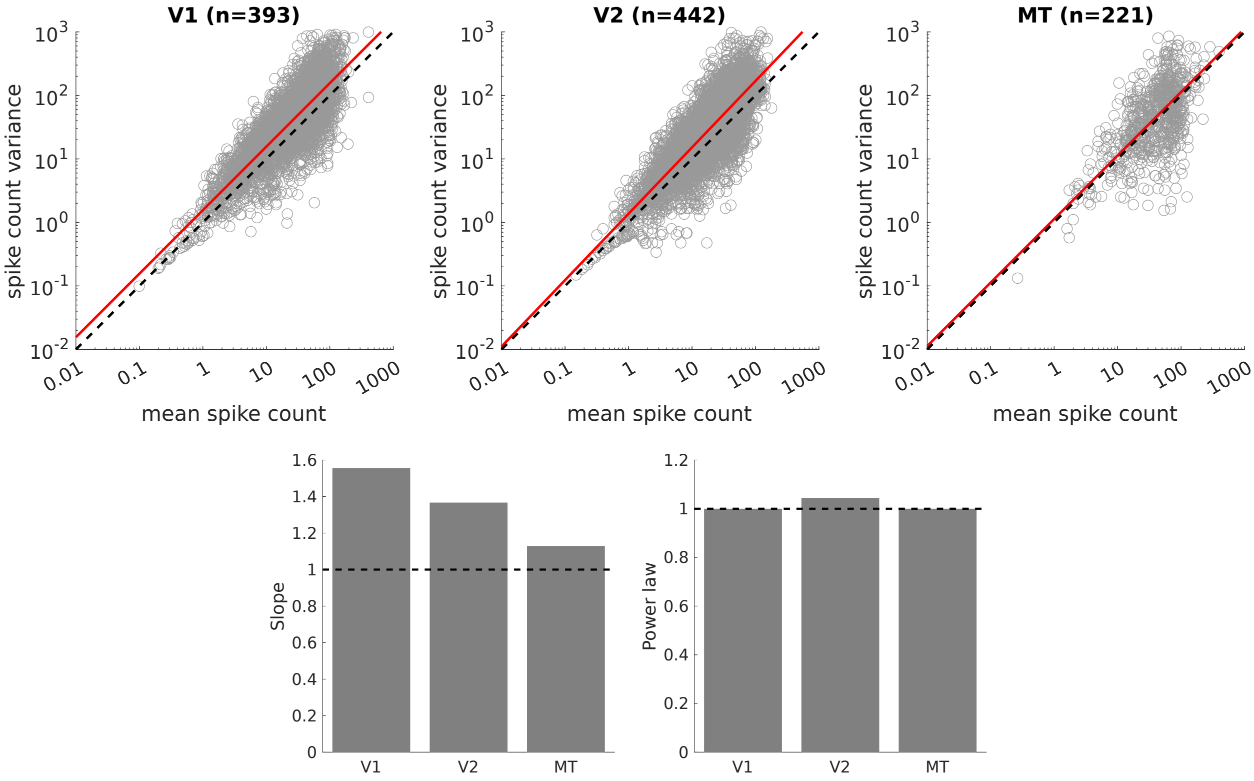


**Fig. S3. (Top)** Relationship between spike count mean and variance for each unique combination of stimulus disparity and cell in each of the three neuronal data sets. The cell sample numbers in the titles reflect the number of cells that contributed to each plot, but each cell contributes more than one data point. Dashed line: unity. Red line: best fit power law function to the distribution. **(Bottom)** The best fit slopes and power laws from each of the red lines in the top set of plots. Dashed line: the expected values of a set of Poisson-distributed spike counts.

### FI distribution divergence using cell count normalization

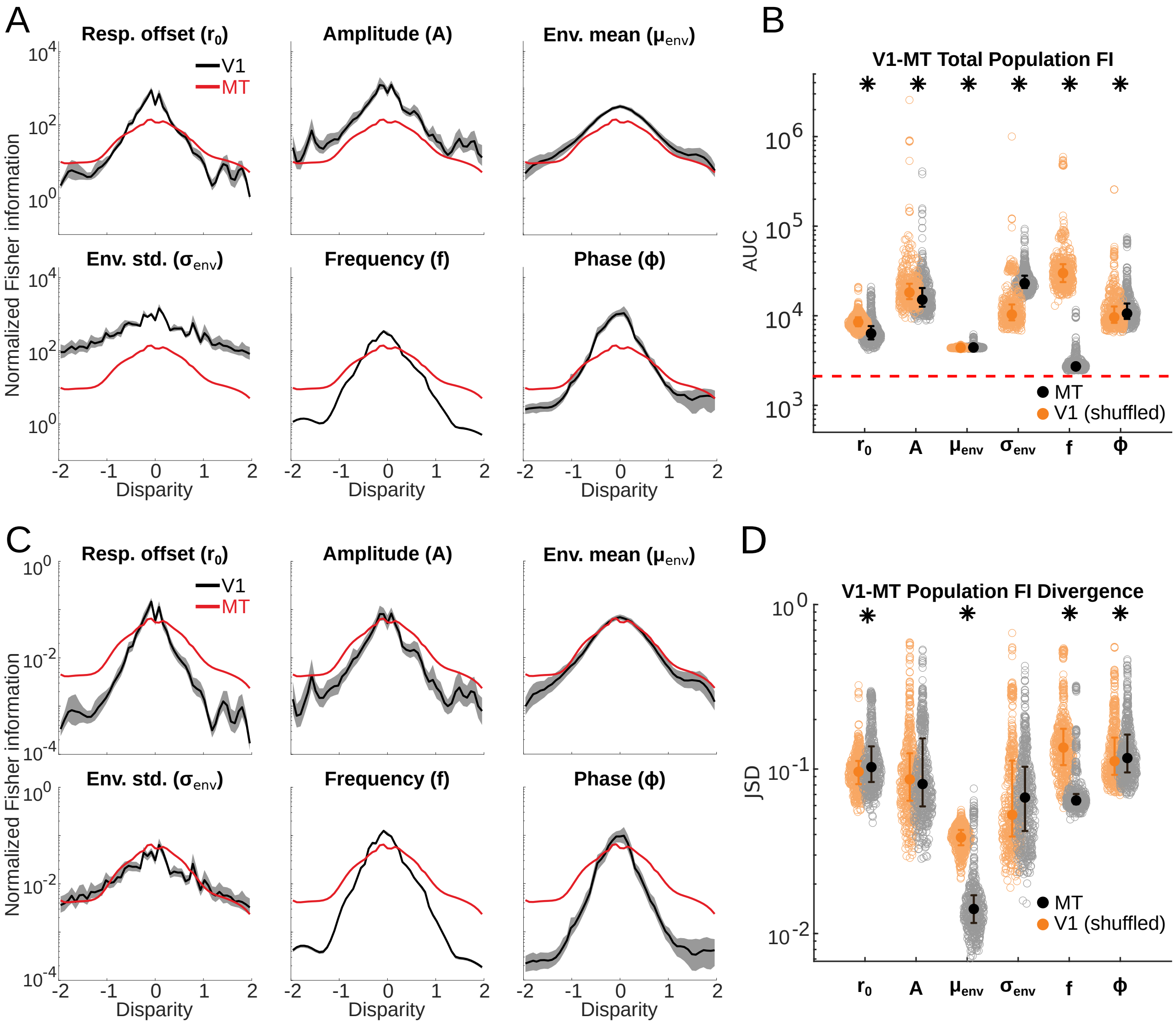


**Fig. S4. A.** Mean and interquartile range from the resampled V1 samples (solid black) compared to the true MT population $FI$ (solid red). Note that the vertical axis is represented on a log scale that exaggerates the difference between distributions at lower probability values. **B.** (black) Area under curve (AUC) reflecting total population Fisher information (TFI) of each bootstrapped V1 population after the set of values for a single parameter are replaced by a set of new parameters sampled from the MT parameter distribution and the empirical MT parameter distribution (dashed red line). (orange) AUC of each bootstrapped V1 population after the set of values for a single parameter are replaced by a set of values sampled with replacement from the V1 population (i.e. the parameters were shuffled between cells). **C.** As in Fig. 5B, but the vertical axis is placed on a log scale for comarison with **A.**. **D.** (black) The same resampled data presented in Fig. 5C (black) JSD when parameters were sampled from the MT parameter distributions and (orange) JSD when parameters were sampled from the V1 parameter distributions (i.e. shuffled). Asterisks indicate significant differences between the two data sets resampled from V1 and MT (full statistics from **B.** reported in Table S4 and from **D.** in Table S5).

### Omnibus tests for inter-area differences in Gabor fits

| parameter | statistics |
|---|---|
| $r_0$ | $\chi^2 = 177$; $df$=2; **p=4.28E-39** |
| $A$ | $\chi^2 = 0.325$; $df$=2; p=0.850 |
| $\mu_{env}$ | $\chi^2 = 11.0$; $df$=2; **p=2.95E-6** |
| $\sigma_{env}$ | $\chi^2 = 251$; $df$=2; **p=3.02E-55** |
| $f$ | $\chi^2 = 47.8$; $df$=2; **p=4.07E-11** |
| $\phi$ | $\chi^2 = 21.2$; $df$=2; **p=2.45E-05** |
| RF center Ecc. | $\chi^2 = 96.9$; $df$=2; **p=8.92E-22** |
| Pref. Disp. | $\chi^2 = 5.16$; $df$=2; p=0.0760 |

**Table S1.** Test statistics and significance for Kruskal-Wallis one-way test for differences between distribution medians comparing the six Gabor parameters across V1, V2, and MT (distributions shown in Fig. 4B). Bolding indicates a statistically significant different between the brain areas, with a significance threshold of $p < 0.05$.

### Paired comparisons of Gabor fits between cortical areas

| parameter | V1-V2 | V1-MT | V2-MT |
|---|---|---|---|
| $r_0$ | z=7.93; **p=1.28E-15** | z=-5.98; **p=2.22E-9** | z=-13.0; **p=1.18E-38** |
| $A$ | N/A | N/A | N/A |
| $\mu_{env}$ | z=-1.27; p=0.202 | z=-4.80; **p=1.57E-6** | z=-4.14; **p=3.42E-5** |
| $\sigma_{env}$ | z=-4.83; **p=1.34E-6** | z=-14.8; **p=1.57E-49** | z=-13.0; **p=1.27E-38** |
| $f$ | z=2.94; **p=0.00325** | z=6.63; **p=3.43E-11** | z=4.98; **p=6.48E-7** |
| $\phi$ | z=2.48; **p=0.0132** | z=-2.44; **p=0.0148** | z=-4.53; **p=5.82E-6** |
| RF center Ecc. | z=-1.87; p=0.0617 | z=-9.03; **p=1.66E-19** | z=-8.64; **p=5.40E-18** |
| Pref. Disp. | N/A | N/A | N/A |

**Table S2.** Test statistics and significance for two-tailed Wilcoxon rank sum test for differences between distribution medians, used for pair-wise follow up tests for the Kruskal-Wallis tests from Table S1. Pairs of distributions tested are indicated in the top row.Bolding indicates a statistically significant different between the brain areas, with a significance threshold of $p < 0.05$.

### Resampled V1 parameter comparisons (Jensen-Shannon divergence)

| | $r_0$ | $A$ | $\phi$ | $\sigma_{env}$ | $f$ | $\mu_{env}$ |
|---|---|---|---|---|---|---|
| $r_0$ | - | z=5.87; **p=4.26E-9** | z=-7.80; **p=6.40E-15** | z=14.1; **p=3.60E-45** | z=19.0; **p=4.12E-80** | z=27.2; **p=2.85E-163** |
| $A$ | - | - | z=-10.4; **p=1.67E-25** | z=8.94; **p=4.06E-19** | z=7.37; **p=1.67E-13** | z=26.5; **p=5.01E-155** |
| $\phi$ | - | - | - | z=-17.7; **p=3.70E-70** | z=-21.8; **p=2.88E-105** | z=-27.3; **p=1.74E-164** |
| $\sigma_{env}$ | - | - | - | - | z=-3.63; **p=2.89E-4** | z=-25.3; **p=1.66E-141** |
| $f$ | - | - | - | - | - | z=-26.7; **p=1.71E-157** |
| $\mu_{env}$ | - | - | - | - | - | - |

**Table S3.** Test statistics and significance for two-tailed Wilcoxon rank sum tests for differences in Jensen-Shannon Divergence (JSD) between each pair of bootstrapped V1 populations generated by resampling from the MT distribution of a given Gabor parameter (i.e. between sets of gray circles in Fig. 5). Members of each pair tested are indicated by the row and column labels.

### Resampling from MT vs V1 (area under curve)

| parameter | MT vs. V1 resampling |
|---|---|
| $r_0$ | $z = -14.95$; **p=1.56E-50** |
| $A$ | $z = -9.72$; **p=2.60E-22** |
| $\mu_{env}$ | $z = 5.03$; **p=4.98E-7** |
| $\sigma_{env}$ | $z = 19.8$; **p=2.59E-87** |
| $f$ | $z = -27.4$; **p=5.86E-165** |
| $\phi$ | $z = 5.36$; **p=8.42E-8** |

**Table S4.** Test statistics and significance for two-tailed Wilcoxon rank sum test for median total Fisher Information (TFI) differences (i.e. comparing the orange and gray bootstraps in Fig. S4B) between bootstrapped V1 populations resampled from V1 and MT parameter distributions for each resampled parameter.

### Resampling from MT vs V1 (Jensen-Shannon divergence)

| parameter | MT vs. V1 resampling |
|---|---|
| $r_0$ | $z = 3.70$; **p=2.12E-4** |
| $A$ | $z = -1.35$; p=0.177 |
| $\mu_{env}$ | $z = -22.1$; **p=5.01E-108** |
| $\sigma_{env}$ | $z = 0.374$; p=0.708 |
| $f$ | $z = -21.1$; **p=1.84E-98** |
| $\phi$ | $z = 2.85$; **p=4.37E-3** |

**Table S5.** Test statistics and significance for two-tailed Wilcoxon rank sum test for median Jensen-Shannon Divergence (JSD) differences (i.e. comparing the orange and gray bootstraps in Fig. S4D) between bootstrapped V1 populations resampled from V1 and MT parameter distributions for each resampled parameter
